# Rise of the Cybercrabs: how digital cloning in an integrated taxonomic framework can support deep-sea exploration

**DOI:** 10.1101/2023.03.16.532964

**Authors:** Emmanuel G. Reynaud, Luis Gutierrez-Heredia, Amy Garbett, Esben Horn, Jens Carlsson, Patrick C. Collins

## Abstract

Taxonomy has been a labour-intensive field of expertise based on hours of manual work and lengthy comments that are mainly bound to books and publications on a two-dimensional world. But every species described is a three-dimensional organism that needs to be seen, manipulated to be fully understood in its native shape. Nowadays, digital technology allows us to transform everyone in an avatar or a digital clone with ease, but collections do not provide many type specimens in a digital format. Here we present a simple approach used to study a specific deep-sea crab *Segonzacia mesatlantica* and provide online digital taxonomy across four repository sites. This offers the possibility to describe, exchange digitally and analyse specimens in its full 3D, establish their taxonomy and share them widely on online databases as well as physically by additive manufacturing to duplicate them in collections and outreach activities. Using an integrated taxonomic approach that included the use of 3D type specimens and molecular barcoding we provide evidence that the genus *Segonzacia* may be more diverse than previously understood

## INTRODUCTION

Biodiversity studies in the deep-sea are inherently difficult and data poor due low spatial coverage and low sample numbers. The high expenses associated with sampling expeditions oftentimes necessitates international collaborations (e.g., Collins *et al*., 2020). This can lead to a sparse and global dispersal of type specimens, especially when the study sites are outside national jurisdictions or in states with a low capacity for specimen curation (e.g., Collins *et al*., 2013, Boschen *et al*., 2015). The realisation of taxonomic studies is therefore, often compounded by the high costs associated with the requirement for intercontinental travel for systematic revisions or inherently risky shipments of type specimens in between collections.

Good taxonomic resolution is fundamental for deep-sea environmental management to provide the base units for statistical analyses and the establishment of conservation objectives (Collins *et al*., 2013). This is critical for low-data ecosystems targeted for marine mining, especially deep-sea seafloor massive sulphides, manganese nodules and cobalt crust deposits (Van Dover *et al*., 2020). The high cost and low throughput of existing taxonomic tools for establishing baselines for deep-sea ecosystems management has been flagged as a concern (Glover *et al*., 2018).

Genetic approaches, such as the Barcode of Life programme, have improved the speed, precision, and accuracy at with operational taxonomic units (OTU) can be delimited and described (Hebert *et al*., 2003). However, the reliability of molecular barcodes to identify OTU’s is hindered by heteroplasmy, mitochondrial introgression, hybridisation, homoplasy, and homologs, (e.g., Magnacca & Brown, 2010). Classical taxonomy’s reliance on high-competence approaches such as line drawings and dichotomous keys is a significant bottleneck leading to a “*gap in scalability*” with species being delimited genetically but lacking a morphological description (Faulwetter *et al*., 2014). Given the low priority to train morphological taxonomists and the increasing power of molecular approaches the disparity between the amounts of data generated relative to the taxonomic output is likely to increase further. An integrated taxonomic approach incorporating multiple approaches, including morphological, is preferable for OTU classification and description (Dayrat, 2005; Will *et al*., 2005).

Digitised and online reference collections may address some of the scalability gap (Godfray, 2007). However, existing online taxonomic image-based tools do not account for morphological variability and are not suitable for systematic revisions (e.g. <u>Mollusca Type Catalogue</u>, <u>Marine Species Identification Portal</u>). Non-destructive 3D imaging techniques, such as CT scanning or photogrammetry, that allow for the detailed reconstruction of the morphology of specimens may support systematic digital revisions (Akkari *et al*., 2015). Cybertaxonomy, a contraction of cyber-enabled taxonomy, is an integrative approach that shares the same traditional goals as taxonomy. Through the adoption of digital technologies and the use of digital platforms, cybertaxonomy can produce results faster and better than ever before (Wheeler 2008; 2010). Existing low cost, rapid and scalable means of sampling and *in situ* digitisation can generate coherent 3D models with high resolution features and mapping competent for taxonomic description and comparative analyses (Gutiérrez-Heredia *et al*., 2015; Gutiérrez-Heredia *et al*., 2016). These low-cost 3D imaging technologies are complemented with digital processing to create 3D datasets of museum collections at different scales (cm to mm) (Nguyen *et al*., 2014, Niven *et al*., 2009).

The concept of using 3D imaging to set up frameworks for creating interactive digital collections and libraries is not recent (e.g., Becerra, 2003). Some of the key elements in the development of this framework included imaging of fine morphological details, accurate 3D model quantification of physical, 3D anatomical reconstructions of soft and hard tissues, complementation of imaging techniques and the creation of online databases with worldwide access (Klaus *et al*., 2003; Raz Bahat *et al*., 2009; Berquist *et al*., 2012). Most studies have relied on expensive and non-portable approaches such as computerized axial tomography scanners, magnetic resonance imaging – which have further limitations in terms of specimen comparisons (Hipsley *et al*., 2020; Morris *et al*., 2018). Recently, 3D datasets of specimens were generated economically and quickly by photogrammetry, but this material either complemented the holotype specimens (Nguyen *et al*., 2014) or was more artistic in merit (e.g. http://digitallife3d.org). Photogrammetry has many advantages for 3D imaging – it is cheap (supporting citizen science), has existing online infrastructure (e.g. https://sketchfab.com) and processing algorithms are open source (e.g. Wu 2011).

The genus *Segonzacia* (Guinot, 1989) is currently regarded as monotypic, with *S. mesatlantica* (Williams, 1988), a vent endemic crab encountered on numerous fields along the Northern Mid-Atlantic ridge (MAR), South of the Azores, i.e., from N to S, Menez Gwen (850 m), Lucky Strike (1710 m), Rainbow (2251 m), Broken Spur (3350 m) TAG (3650 m), Snake Pit (3480 m), Logatchev (3080 m), Ashadze (4080 m), in populations of low density, on walls of black smokers, associated with assemblages of deep-sea mussels (Williams, 1988; Guinot 1989; 2006; Fabri *et al*. 2011, Mateos *et al*. 2012). The 45° North vent field is north of the Azores and is the only high temperature hydrothermal vent known between the Azores and Iceland. This vent field has a unique geological setting; it is the only pure basalt-hosted tectonically controlled vent site and the only site situated on the median valley wall (Wheeler *et al*., 2013). This isolation may support endemic taxa. However, some studies have demonstrated that certain taxa (e.g., *Peltospira smaragdina*) are found at both the 45° North site and south of the Azores sites (Collins *et al*., 2020).

Here we present a proof-of-concept approach for an integrated taxonomic approach incorporating 5 cybertypes generated by photogrammetry. Virtual volumetric models of deep-sea crabs, recovered from hydrothermal vents along the Mid Atlantic Ridge, were shared online and supported by ecological information, were used to support species identification in a data poor ecosystem. We test, in the absence of conclusive genetic data the question of whether the unique geophysical structure of the Moytirra ecosystem, associated with numerous rare characteristics, hosts a novel species of *Segonzacia*.

## METHODS

### Sampling

Investigations and sampling of the 45° North vent field were carried out on board the research ship RV *Celtic Explorer* and using a remotely operated vehicle (ROV), *Holland 1* during the VENTuRE Survey between 11 July and 4 August 2011. Crab specimens (n = 4) were collected by ROV aspirator (Figure 2A) were provisionally identified as *Segonzacia spp*. (Guinot, 1989; see the key by McLay, 2007). Three crabs were fixed in 5% formalin and preserved in 70% EtOH and one was preserved in 100% molecular grade EtOH for molecular studies. The specimens are deposited with Dr Jens Carlsson Group (Ethanol; University College Dublin, ♂= 2, ♀ =1) and the NUI Galway Museum of Zoology (formalin fixed but now exhibited as a dried specimen, ♂= 1).

These specimens were compared against a confirmed reference specimen of *Segonzacia mesatlantica*, from the Muséum national d’Histoire Naturelle collection, Paris (Ethanol, ♂= 1, MNHN-IU-2014-9439). The reference specimen was collected in 2001 from Menez Gwen vent field. In total we generated 5 digital clones, 4 males including reference specimen and one female (Table I with links to 3D models).

**Table I.** Measurements of digitised specimen files with links to 3D models.

| Specimen | Actual Specimen |  |  | Digital Specimen |  |  |  |
| --- | --- | --- | --- | --- | --- | --- | --- |
|  | Sex | Width (mm) | Length (mm) | Volume (mm<sup>3</sup>) | Vertices | Data size | Link to model |
| TYPE | ♀ | 58.52 | 34.25 | 62188.30 | 384,035 | 213 MB | https://skfb.ly/o66vR |
| Crab A | ♀ | 52.30 | 38.96 | 44404.10 | 805,789 | 322 MB | https://skfb.ly/o66tx |
| Crab B | ♀ | 45.95 | 26.80 | 28495.90 | 1,260,037 | 400 MB | https://skfb.ly/o66vB |
| Crab C | ♀ | 40.89 | 24.24 | 20367.70 | 493,356 | 246 MB | https://skfb.ly/o66tV |
| Crab D | ♀ | 50.92 | 29.95 | 36726.20 | 1,049,599 | 266 MB | https://skfb.ly/o66vE |

### Molecular Barcoding

A crab leg was dissected from a specimen preserved in 100% ETOH (not used for imaging) from the 45° North. Approximately 25mg of tissue was removed from the leg using a sterilised forceps with DNA extracted using the DNEasy Blood and Tissue Kit (Qiagen, Inc. Valencia, CA). DNA concentrations were quality checked using a Nanodrop-1000© (Thermo Fisher Scientific Inc.). The universal COI primers LCO1490 and HCO2198 (Folmer et al, 1994) were used to amplify COI. Mastermixes for PCR were composed of forward and reverse primers, magnesium chloride (MgCl2), deoxyribonucleotides (dNTP’s), double-distilled water (ddH2O), bovine serum albumin (BSA) and Taq polymerase. The final PCR reaction consisted of 23µl of mastermix with 2µl of DNA. To act as a negative control, 2µl of dH2O was added to a tube after every sample. The samples were put in the PCR machine, T3000 (Biolabo, SA) for three hours and then stored in a refrigerator at 4°C. DNA was purified using ExoSAP prior to Sanger sequencing, with 6µl of mastermix and 22µl of product centrifuged to ensure thorough mixing. Samples were incubated in a thermocycler at 37°C for 15 minutes and then 80°C for a further 15 minutes. Samples were then stored at 4°C. A total volume of 3µl per individual was sent off for Sanger sequencing with Marcrogen (http://www.macrogen.com/eng/, 2014).

### Sequence Editing

Chromatograms were imported to Geneious 7 (Biomatters, NZ). Forward and reverse sequences were aligned and any gaps or single nucleotide polymorphisms (SNPs) as well as poor signal areas at the sequence ends were removed.

### Specimen identification

The aligned sequences were uploaded to BLAST using MEGAX software and the most closely related sequences were identified (Tamura et al, 2011). Sequences for the COI gene of other *Segonzacia* specimens were downloaded from GenBank (n=6 (http://www.ncbi.nlm.nih.gov/genbank/, 2022)). The shore crab, *Carcinus maenas* was chosen as outgroup. Sequences were aligned using MUSCLE on MEGA6 (Hall, 2011).

Aligned data was used to generate a maximum Likelihood (ML) tree using MEGAZ. ML trees were chosen as this method takes probability into account and allows the model to fit the small amount of data provided. The ML method assumes that evolution of lineages and branches is independent and allows for it to go forwards and backwards rather than forcing a result (Hall, 2011). The phylogenetic tree was inferred by Maximum-likelihood (ML) analysis, and a bootstrap consensus tree was constructed (see Supplementary Material Figure S1).

### Haplotype Network Analyses

Haplotype networks of the CO1 gene were constructed in PopART (Leigh & Bryant, 2015) using the TCS method (Clement *et al*., 2002) to infer relationships among the sample sequence and all available COI *Segonzacia* sequences (n=6). The haplotype network was produced to visualise interconnectivity and estimated mutations among sequences and colour coded based on region (Figure 1).

**Figure 1:**
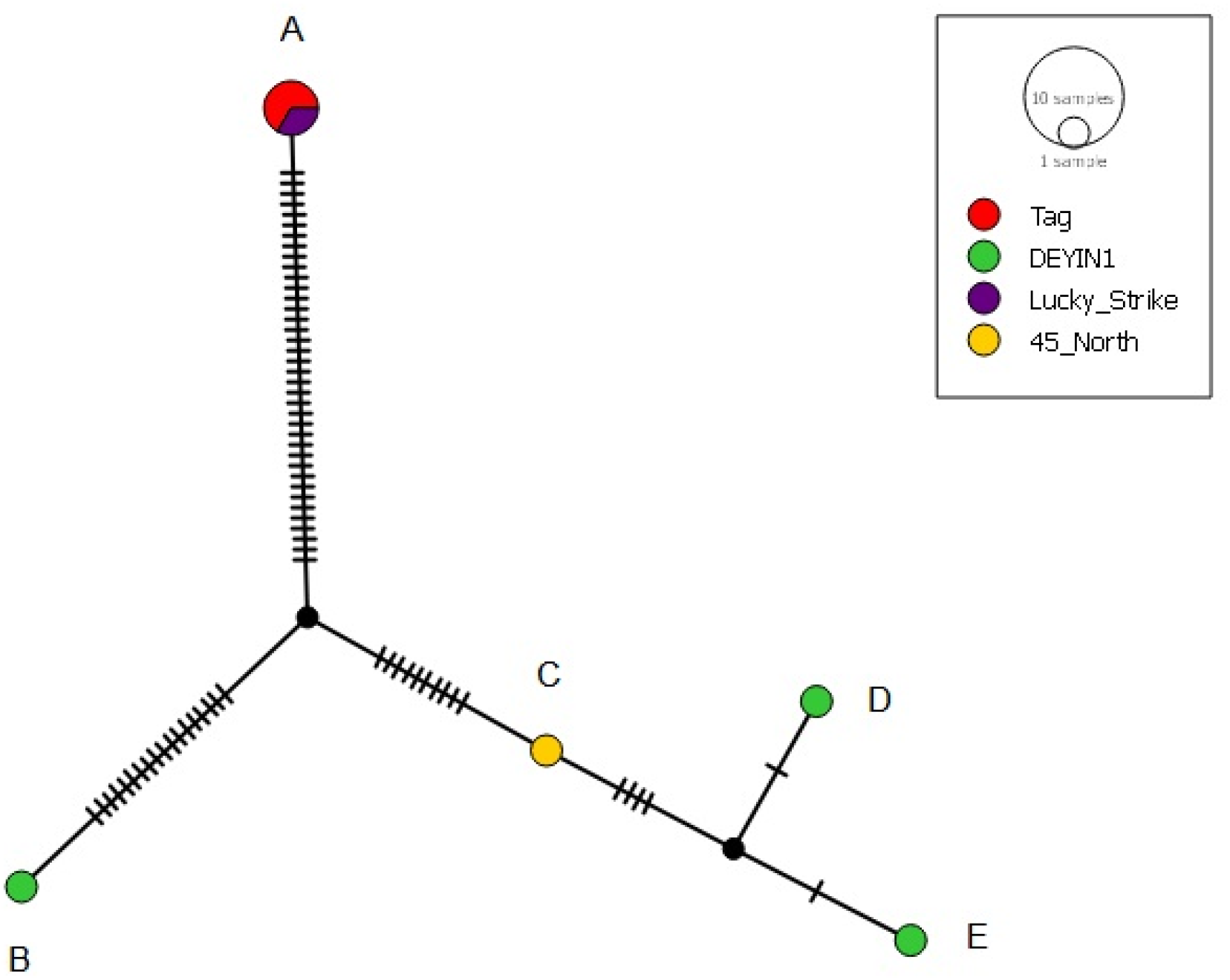
Haplotype network of the CO1 gene for the sample sequence and sequences of *Segonzacia mesatlantica* constructed with the TCS program, unrooted. Each line segment represents an inferred mutation step. Haplotypes are coloured by region and scaled proportionately to the number of supporting sequences. Clusters represent **A)**. *Segonzacia mesatlantica* (NC_035300.1 & KY541839.1 & JQ407473.1) **B)** *Segonzacia mesatlantica* (KY445841.1) **C)** 45 North Sample **D)** *Segonzacia mesatlantica* (KY445843.1) **E)** *Segonzacia mesatlantica* (KY445842.1).

### Image acquisition

Specimens were extracted from their storage vial and allowed to dry to avoid reflection, except for the dry preserved specimen (Galway). Imaging was performed as fast as possible (<30 mins) to avoid damage to the specimen. Images taken using a tripod mounted APS-C sensor digital SLR camera with an attached 100mm f/2.8 Macro lens (TTL metering, @f11). Each specimen was suspended by a clear nylon monofilament with at least two support-points in contact with a patterned stage to minimize movement (Figure 2B). The patterned stage enhances feature recognition and matching over a homogeneous surface. A high-contrast plastic ruler was positioned adjacent to the specimen for scale. Three sets of overlapping images (n = 40) were taken at elevations of 45°, 0° and 215° from the midpoint of the specimen - ensuring 360° coverage of the specimen at each elevation (Figure 2C). A final set of close-up images was taken of taxonomic-diagnostic characterizing features as determined by reference keys and literature to facilitate accurate identification. The final render (see below) is presented in Figure 2D.

**Figure 2:**
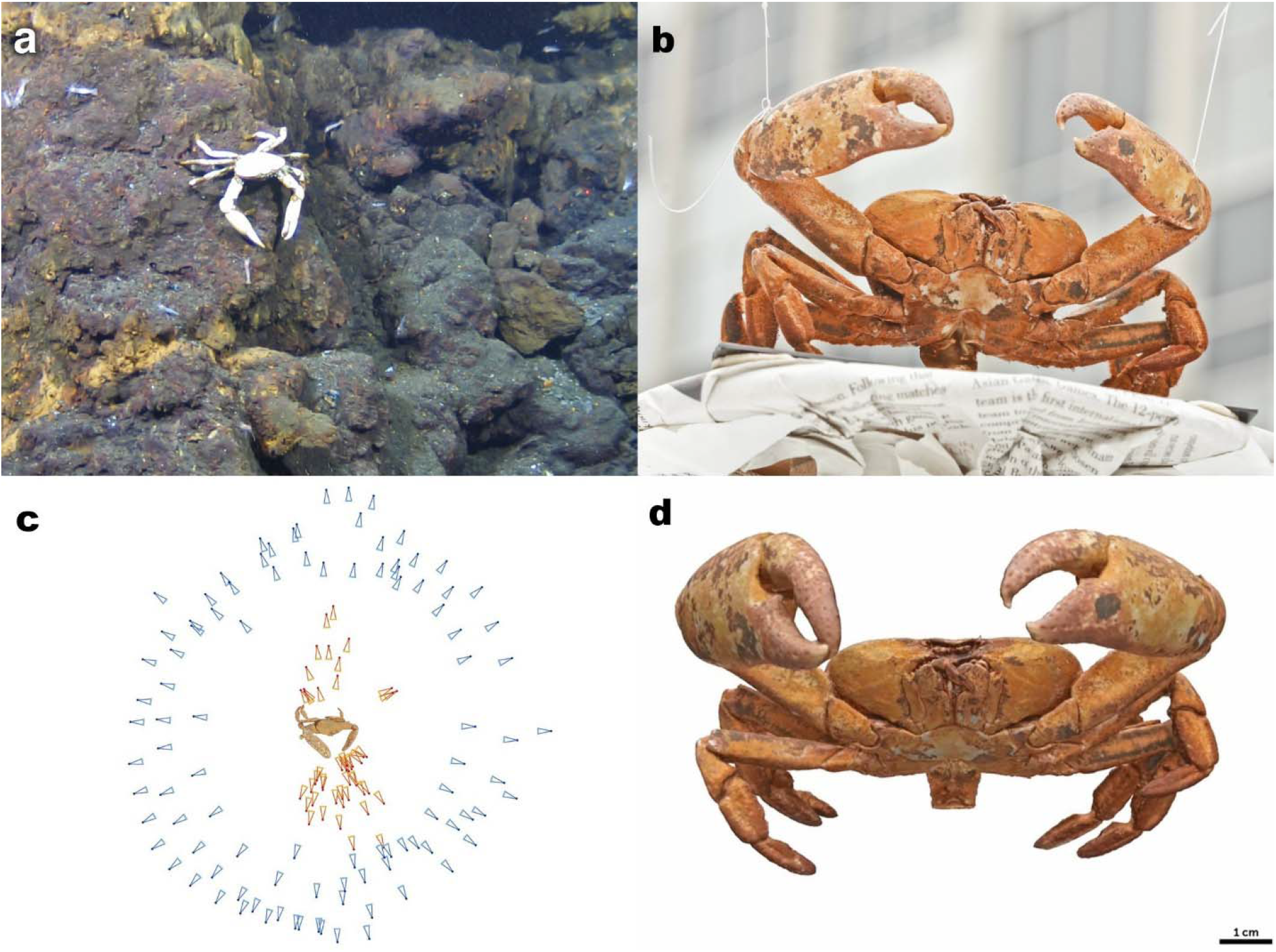
Photogrammetry Workflow. Figure outline workflow for photogrammetry. **a**. live crab institute at the 45° North vent site. **b**. preserved crab positioned for imaging with clear nylon monofilament. **c**. Typical camera angles used to create model. **d**. Screengrab of resulting 3D volumetric model

### Image Processing

All images were batch-edited, maintaining image metadata, using the contrast enhance, light curve correction and image sharpening subroutines in Photoshop CS5 Bridge (Adobe Systems, U.S.A.). Dense reconstruction of image files was carried out using VisualSFM (Wu, 2013). The CMPMVS subroutine of Visual SFM was used for the mesh reconstruction of the dense model. This resulting 3D mesh model was saved as a Wavefront object (.obj file). The models were further cleaned using: Meshlab (Cignoni *et al*., 2008) for scale correction using a reference rule; Meshmixer for hole repair and mesh cleaning (www.meshixer.com). The model processing pipeline is outlined in Figure 3.

**Figure 3:**
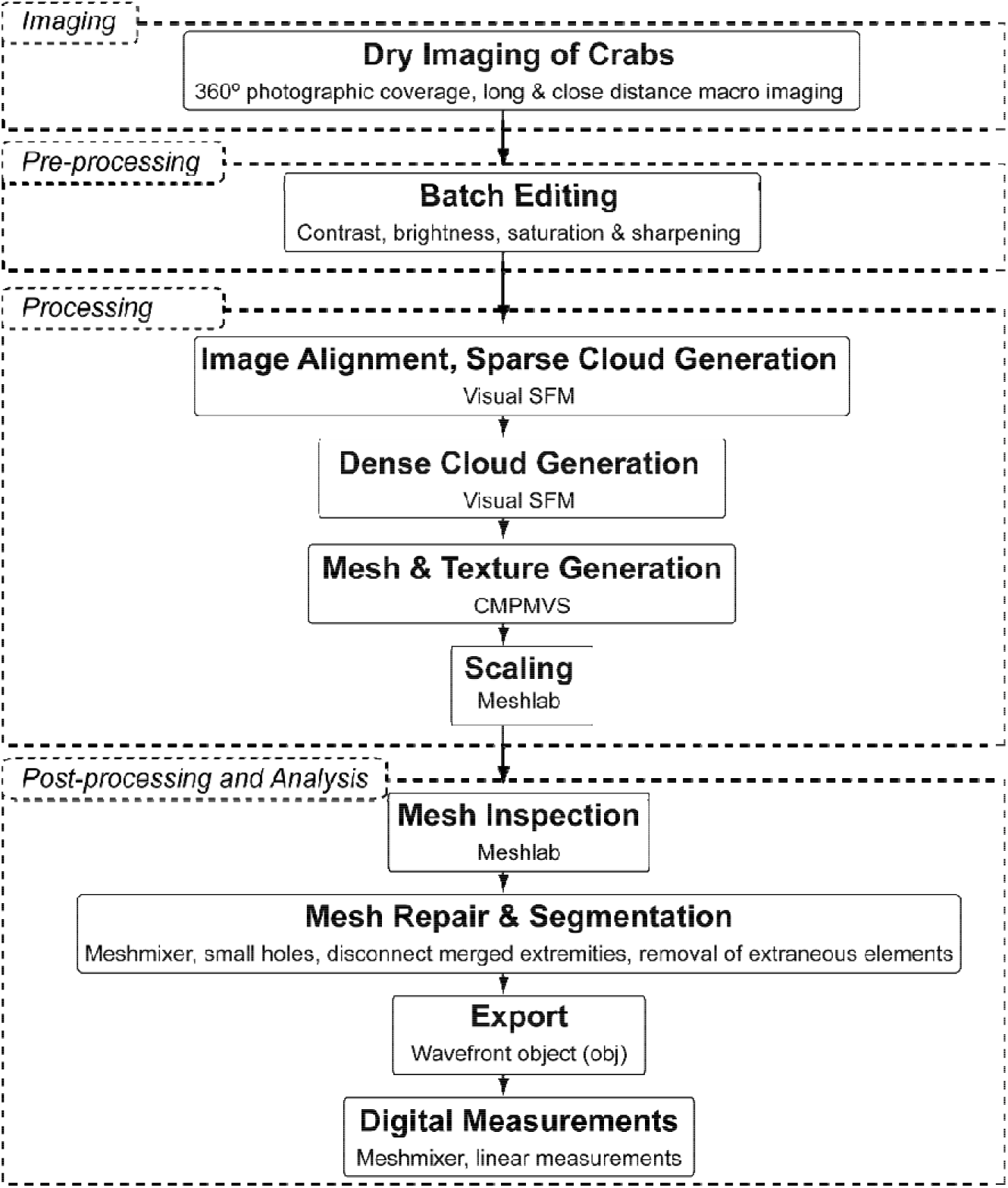
A flowchart highlighting the image processing pipeline.

### Morphometric analysis

Morphometrics analysis to compare specimen morphology and establish their variability was performed using Open Source CloudCompare platform [http://www.cloudcompare.org/]. In order to perform a meaningful comparison, we segmented the shell of each crab including point cloud and mesh map. They were then imported into the CLoudCompare platform in pairs, followed by transformation, registration to align both shells to allow for point cloud to mesh comparison. This allows for a complete 3D morphometric analysis by visualization of colour depth between the reference shell as chosen by the user.

### Validation and demonstration of cyber-specimens

Images (2D) of live crab *in situ* were shared digitally, however this was not sufficient to decide of the species was different from *S. mesatlantica*. Therefore, 3D model crabs (cyber-specimens) of the 45° North vent field were shared digitally demonstrate the applicability of the approach for taxa identification. Furthermore, cyber specimens were shared with the artist Esben Horn (10TONS, Denmark) for duplication using 3D additive manufacturing and artistic colouring to test the possibility to duplicate the model from physical to digital and back for outreach, display, and comparative analysis.

## RESULTS & DISCUSSION

Molecular analyses (COI) of the 45° North vent field crab specimens identified *Segonzacia mesatlantica* as the closest match. However, the genetic evidence was not equivocal (see Figure 1 and supplementary material Figure 1) due to the small sample size of specimens, it suggested that the specimens were at least in the *Segonzacia* genus. Phylogenetic and haplotype analysis indicated the 45° North sample were comparable to *Segonzacia mesatlantica* (Accession numbers KY445843.1 & KY445842.1) from the Deyin1 region. However, the results indicate this three clades, Tag and lucky strike region, Deyin 1 and a clade with both some Deyin 1 and 45° North samples (see Figure 1 and supplementary material Figure 1). Evidence here suggests novel species of *Segonzacia*, with the 45° North sample and Deyin of *Segonzacia mesatlantica* being distinct from the Tag and Lucky Strike specimens of *Segonzacia mesatlantica*. While the molecular evidence is not sufficient to support the thesis that the Moytirra vent field hosts unique fauna, it did extend our understanding of the phenotypic variation found in *Segonzacia* (with perhaps more species diversity).

Comparisons made using the 3D imaging highlighted that the chelipeds of *the* 45° North *Segonzacia* spp. appeared less massive with a more pronounced curvature and more highly tapered dactylus tips when compared to the reference *Segonzacia mesatlantica* specimen (Figure 4) from the Menez Gwen site. These are likely adaptations to the Moytirra prey choice (*sensu* Lee & Seed, 1992). The most obvious food source at the 45° North vent site were the thumbnail-sized limpet, *P. smaragdina*, with densities of up to 50,000/m^2^ (Collins *et al*., 2020). This is unlike the south of the Azores vent sites where the larger and more robust *Bathymodiolus* spp. mussels are likely the primary food source (Cosel & Comtet 1999, Cosel *et al*., 1994). This may be evidence of some niche separation among *Segonzacia* crabs. Slight differences were also observed in relation between the 45° North specimens and the type specimen (holotype) female and the gonopods of male *Segonzacia mesatlantica* specimen described by Guinot (1989). In the type specimen (Guinot, 1989), the G1 is nearly straight, thick, with a wide, bilobate apex, bearing long setae. The G2 have the two long flagella that interlink and are corkscrew-shaped, the end of the spiral hardly extending beyond the G1 tip (but, once uncoiled, the total length of the spiral is actually greater than the G1). The ensemble G1+G2 is proportionally short, reaching the thoracic sternal suture 5/6 (setae not counted). In the 45° North specimen*s* (Crab A, Figure 4) the G1 and G2 are roughly similar in shape, but the G1 is significantly arched, forming a strong curve, with the apex and the setae; the G2 do not interlink, have a straight, longer flagellum that ends in a short, coiled part, reaching thoracic suture. However, the resolution of the digital clones was not sufficient to fully resolve these differences and required additional imaging, highlighting a shortcoming in the approach (Figure 5). It should be noted that the gonopod structure should not be considered taxonomically diagnostic in this situation due to the low number of specimens observed. However, the lines of evidence from an integrated taxonomic approach incorporating both molecular barcoding and the 3D model comparison suggest that the 45° North *Segonzacia* are ecologically and evolutionary distinct from *Segonzacia mesatlantica* and more like some crabs found at Deyin 1.

**Figure 4:**
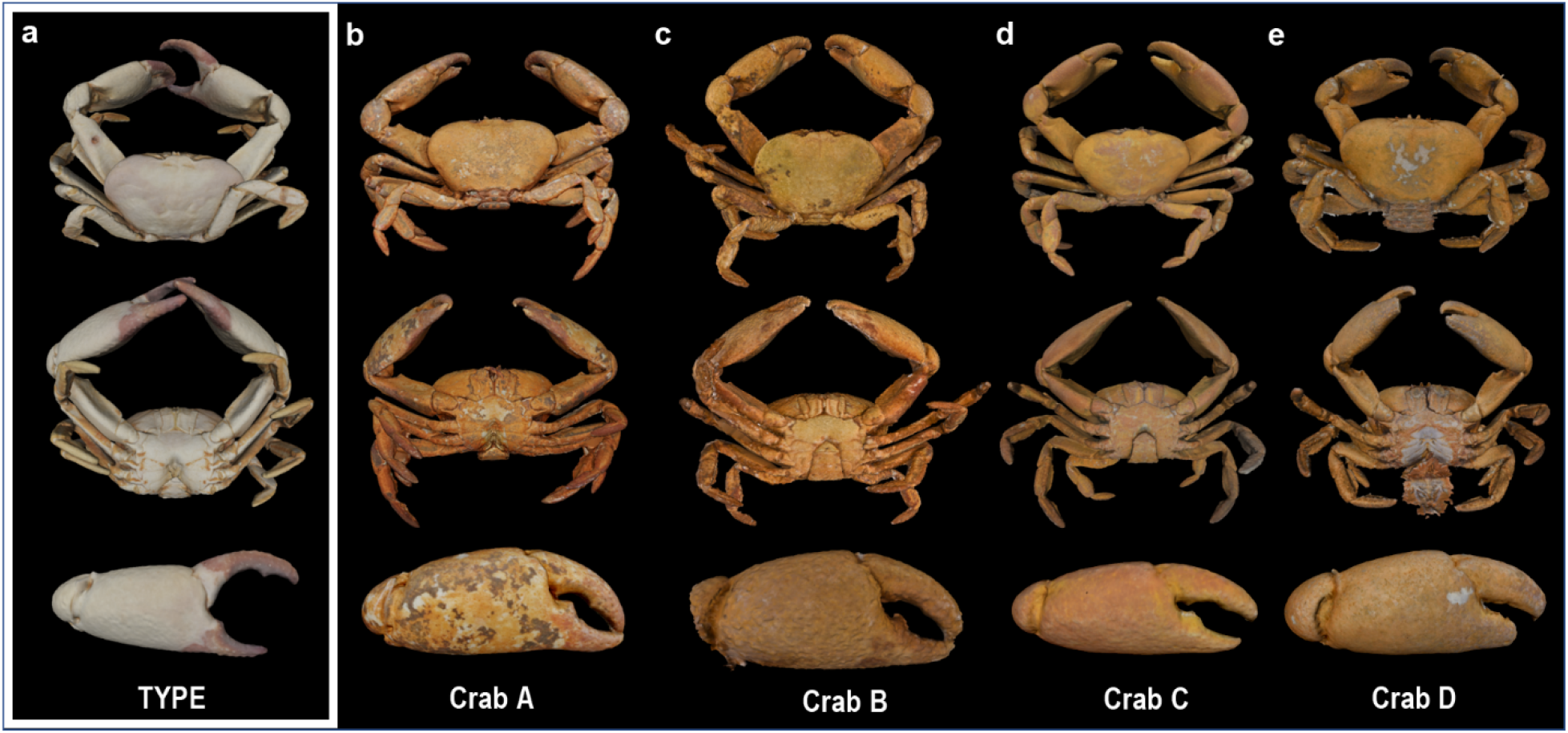
*Segonzacia mesatlantica* digital clones. **a**. Reference male specimen from the Museum national d’Histoire Naturelle, Paris. **b**. Ethanol preserved male specimen from 450N. **c**. Dry preserved male specimen from 450 N **d**. Ethanol preserved male specimen from 450 N **e**. Ethanol preserved female specimen from 450N. Top row highlights the dorsal view; middle row highlights the ventral view and bottom row highlights the left cheliped. All models are scaled to the same size for ease of comparison (measurements are presented in Table I)

**Figure 5:**
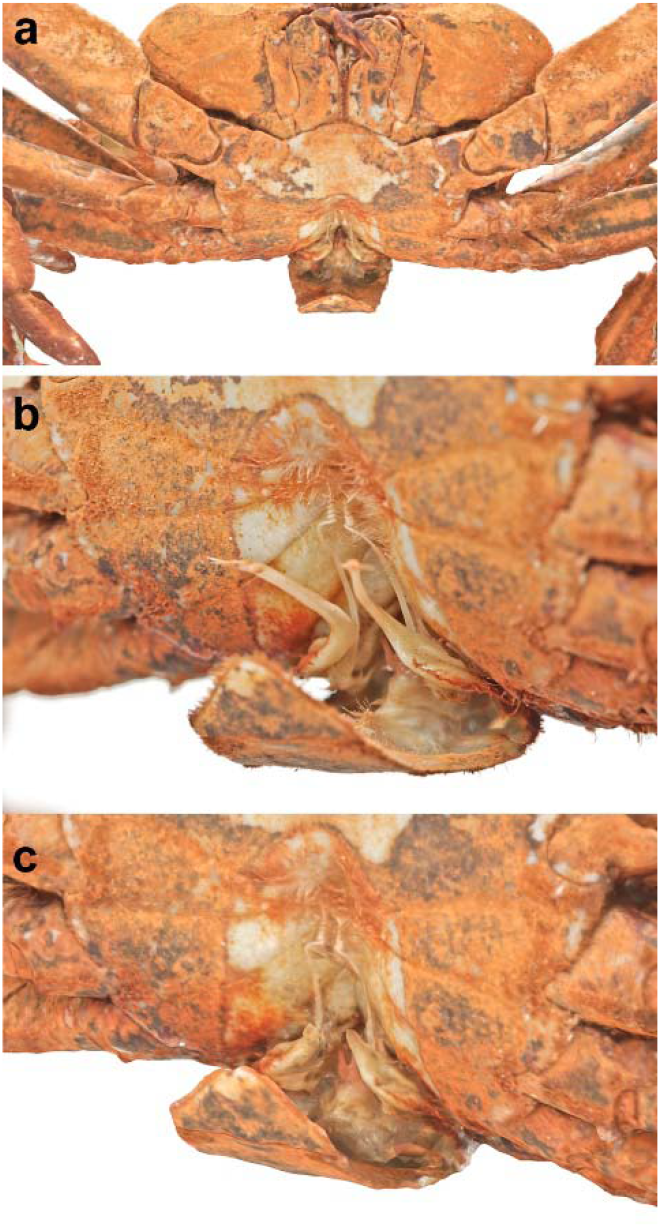
Photogrammetry limitations. **a**. Front view of the male specimen from 450N (crab A in Figure). **b**. Original non-processed close-up image highlighting sexual appendages **c**. Processed close-up of the 3D digital model highlighting poor resolution.

The photogrammetry imaging succeeded in producing accurate 3D digital clones that displayed visually satisfying levels of morphological detail in terms of texture, colouration, and geometry (Figure 3, Sketchfab links in Table I). That all four crabs could be confirmed as *Segonzacia* using only digital clones validates the applicability of photogrammetry for deep-sea digital taxonomy. Furthermore, the technique is cheap and can be performed nearly anywhere and preserved the texture and colour of the original specimen normally lost during CT scans. However, considerable knowledge gaps remain, with only morphological and molecular data existing for the 45° North samples. More samples are required to confirm the presence of a second vent crab on the mid Atlantic Ridge, *Segonzacia cf. cybertaxa*.

The study did highlight the efficacy of 3D imaging for deep-sea taxonomic studies. Probably the most pertinent application of the digital clones is allowing for the sharing of samples safely without the need of CITES permit across the globe in a manner that allows for online comparison but also digital replication using 3D printers. This is an attractive approach that can reduce shipping costs of specimens for the hosting institution. In the current pandemic climate, this allows for the continuation of vital collaborations between taxonomists. The development of taxonomic monographs usually requires either the chaperoning of samples by scientists to places of research or the travel of scientists to institutes hosting specimens. These activities require the movement of either scientists or specimens which can constitute a real risk to both, especially to scientists during a global pandemic.

Another attractive aspect of generating 3D models is the possibility to create automated full 3D morphometric digital analysis as currently used in engineering and industry related standards. Point cloud to mesh comparison (Figure 6) displays a quantitative visualization of errors and standard deviation of topology related structures. This also permits the creation of reference atlases to average population variation. Moreover, this is a fully quantitative approach that can lead to automated recognition and ecological analysis whatever the initial position of the picture taken for in-field identification or citizen-science queries. Those 3D morphological atlases can be used for population studies as they are a digital reference type.

**Figure 6:**
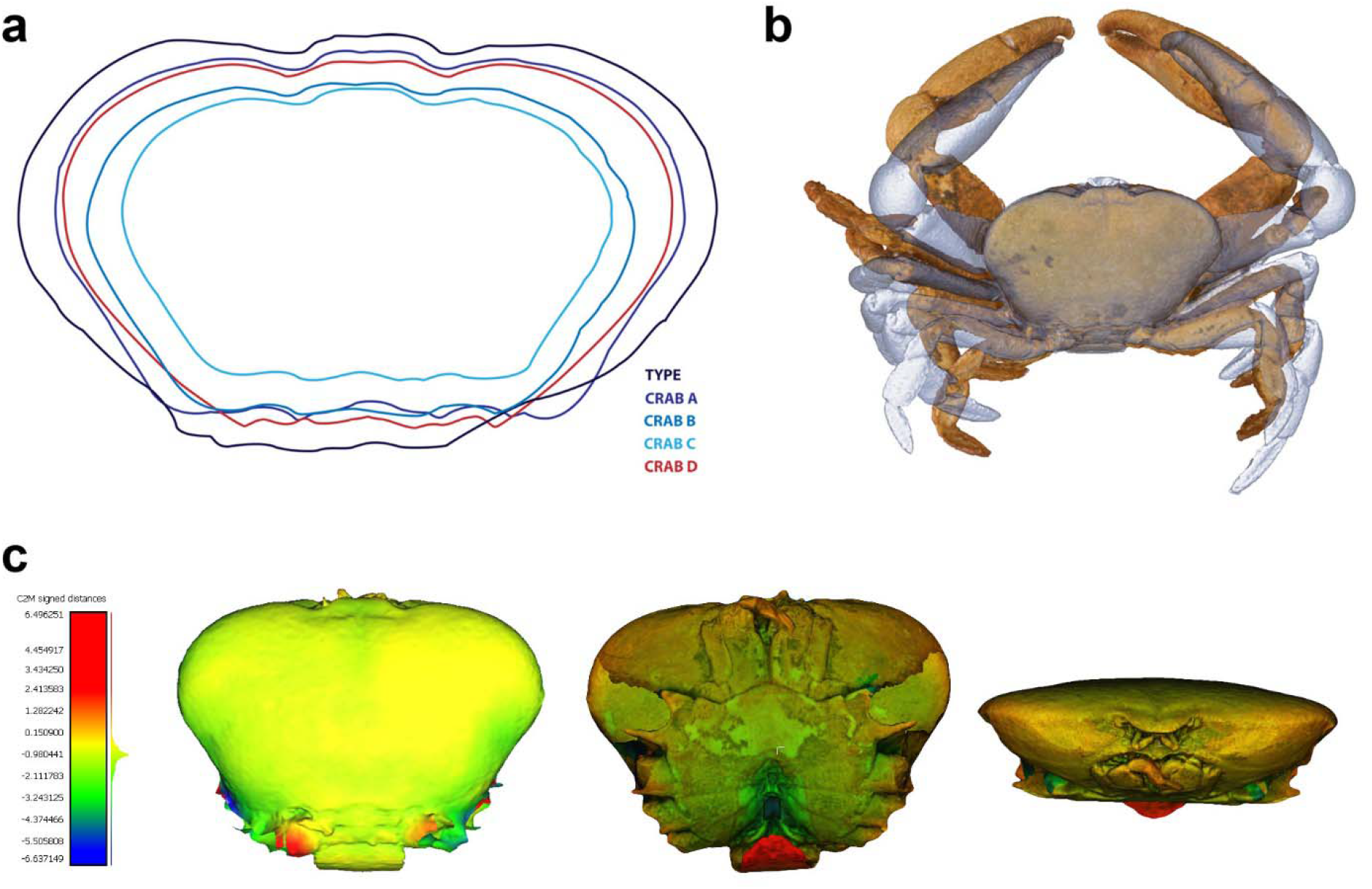
Morphometric analyses of specimens. **a**. Classical alignment of shell contours extracted from the 3D models used for PCA analysis. **b**. Attempt to align the complete model of 2 crabs limited by their initial position. **c**. Point cloud to mesh comparison of Menez Gwen specimen specimen and Crab A from the Moytirra site. Green to red colour relate to alignment quality from best to worst.

The application of cybertypes, not based on the type, goes beyond just digital-holotyping. The data from multiple specimens, when combined as an atlas, can act as reference dataset that incorporates an element of variability (Baldock *et al*., 2001). Artificial intelligence-based approaches can even recognise portions of a specimen and identify to species independent of pose (e.g. Nguyen *et al*., 2017). The variability inherent in an atlas-based approach can support both the identification of species and allows for phylogenetic analyses. Furthermore, with more data, this approach could support atlas-based population assessments – phenotypic expression of the genome. This could potentially allow for rapid population assessments without hard sampling. This would be especially useful for high conservation survey, for example at deep-sea hydrothermal vents. This would support linkage between the genetic and morphological databases. Such linkage would improve quality control of molecular reference collections (e.g.Genbank) as they would have a secondary point of comparison.

Identification using convolutional neural networks (*sensu* Miao *et al*., 2019) has been demonstrated successfully for the identification of fish species (Iqbal *et al*., 2019), freshwater invertebrates macrofauna species (Ärje *et al*., 2020) tiger individuals (Shi *et al*., 2020) and even diatoms (Kloster *et al*., 2020). Such approaches can support citizen science based taxonomic studies and allow for the scientific interrogation of industrial exploration footage (e.g., ROV data from subsea pipeline or mining programs). The creation and sharing of 3D digital holotypes (e.g., using Sketchfab or Morphosource) allows educators, digital content creators and artists access to museum quality specimens. This can support wider scientific outreach and increase public awareness of otherwise poorly known ecosystems (e.g., in the deep sea.

The adoption of digital cybertyping can lower the carbon footprint of researchers and increase the security of priceless type specimens. Neither scientists nor specimens need to travel for research to be undertaken, digitised specimens can be shared via the web. This can support open science, allowing for data mining at a larger scale than currently feasible. The digital clones can support educational outreach; 3D printing can allow for rare specimens, especially relevant for deep-sea, to be shared between museums and educational institutes (Figure 7).

**Figure 7:**
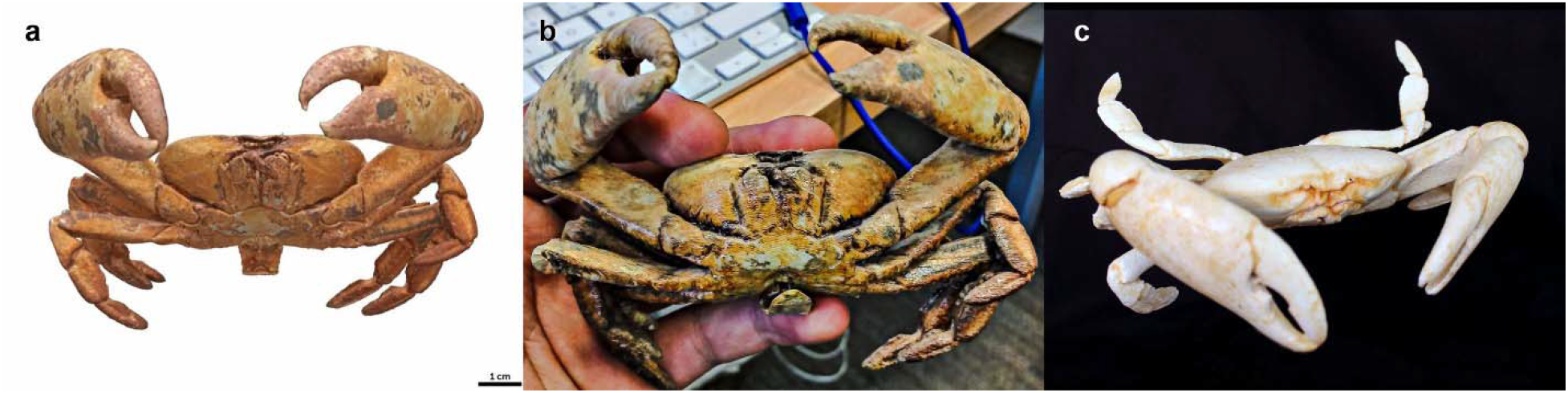
Outreach use of 3D digital models. **a**. a still image of a specimen (see video in supplementary material) **b**. McCor paper 3D printed model for museum display. **c**. 3D printed and painted model generated from files in Copenhagen.

## ACKNOWLEDGEMENTS

We thank the Muséum national d’Histoire Naturelle, Paris, that provided access to the reference specimen [MNHN-IU-2014-9439 = MNHN-B28370]and to Prof. Danièle Guinot who examined a specimen from the Moytirra site in 2015 and shared her observations with us. We acknowledge the support from Science Foundation Ireland grant 12/IP/1308. The sampling expedition, VENTuRE survey, was principally funded by the Marine Institute under the 2011 Ship-Time Programme of the National Development Plan and by the National Geographic Society with additional support from National Geographic Television, National Oceanography Centre, UK, University of Southampton, UK, Geological Survey of Ireland (INFOMAR programme), University College Cork and National University of Ireland, Galway.

## LINKS

### Videos link

Celtic Explorer survey 2011 (see 2.21 minutes)

https://www.youtube.com/watch?v=g0O-KmFwKHc

**SketchFab links**

**Segozancia sp. male crab (TYPE)**

https://sketchfab.com/3d-models/segozancia-sp-male-crab-type-53a51432f37142f1a0b86a6883581e58

**Segozancia sp. Paratype (Crab A)**

https://sketchfab.com/3d-models/segozancia-sp-paratype-crab-a-e54d98b7af734c19ab67d8b03ecad5ff

**Segozancia sp. Male crab (Crab B)**

https://sketchfab.com/3d-models/segozancia-sp-male-crab-crab-b-ea0f2a919898477a8de9682f107e181c

**Segozancia sp. male crab (Crab C)**

https://sketchfab.com/3d-models/segozancia-sp-male-crab-crab-c-8137509bf95a4f799d8715d516aa00b8

**Segozancia sp. Female crab (Crab D)**

https://sketchfab.com/3d-models/segozancia-sp-female-crab-crab-d-add59ee18dd74ffa9d1bd5495da3e1f3

## SUPPLEMENTARY MATERIAL

**Figure 1.**
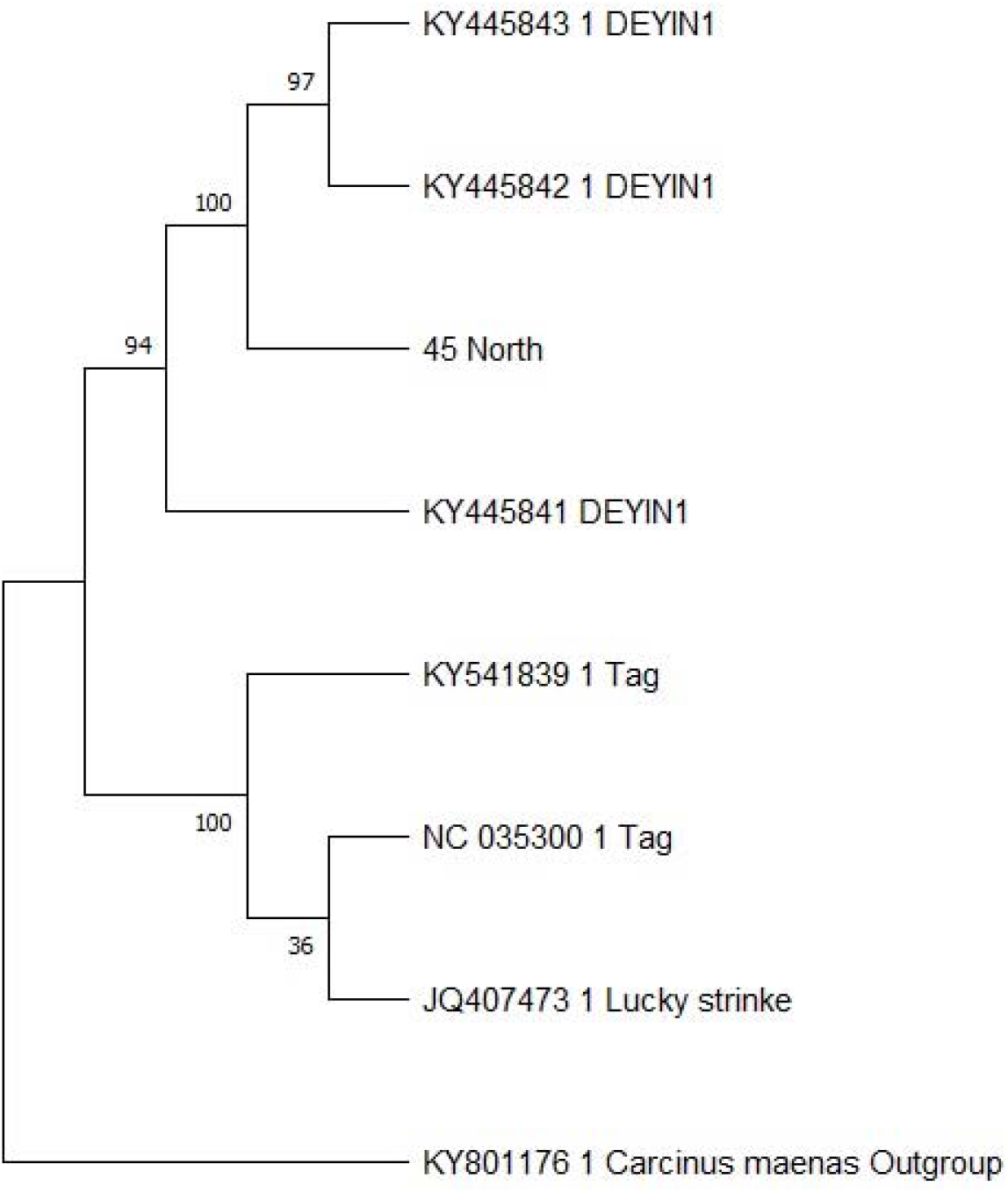
Phylogenetic analysis of *Segonzacia mesatlantica* species. COI phylogenetic gene tree for *Segonzacia mesatlantica* constructed using Maximum Likelihood method, bootstrap consensus tree rooted with the outgroup *Carcinus maenas*.

